# Hippocampal sequences represent working memory and implicit timing

**DOI:** 10.1101/2025.03.17.643736

**Authors:** Conor C. Dorian, Jiannis Taxidis, Dean Buonomano, Peyman Golshani

## Abstract

Working memory (WM) and timing are considered distinct cognitive functions, yet the neural signatures underlying both can be similar. To address the hypothesis that WM and timing may be multiplexed we developed a novel rodent task where 1^st^ odor identity predicts the delay duration. We found that WM performance decreased when delay expectations were violated. Performance was worse for unexpected long delays than for unexpected short delays, suggesting that WM may be tuned to expire in a delay-dependent manner. Calcium imaging of dorsal CA1 neurons revealed odor-specific sequential activity tiling the short and long delays. Neural sequence structure also reflected expectation of the timing of the 2^nd^ odor—i.e., of the expected delay. Consistent with the hypothesis that WM and timing may be multiplexed, our findings suggest that neural sequences in dorsal CA1 may encode cues and cue-specific elapsed time during the delay period of a WM task.

## INTRODUCTION

Working memory (WM) is a fundamental component of cognition. It is generally defined as the ability to temporarily store and flexibly use information to guide goal-oriented behaviors [1, 2]. Early pioneering studies of WM suggested that steady-state persistent activity—fixed-point attractors in the language of dynamical systems—is the dominant neural mechanism underlying WM [3–5]. However, in the past decades, a number of other dynamic regimes have been associated with WM, including neural sequences [6–9] and ramping activity [10–16].

Another critical component of cognition, and behavior in general, is the ability to tell time and anticipate when external events will occur on the scale of seconds [17–20]—the same time scale of working memory. Indeed, even when the timing of stimuli and temporal structure of a task is irrelevant to the task itself, humans will implicitly learn the temporal structure as a means to prioritize attention and optimize cognitive resources [18, 21, 22]. As in the WM field, the neural mechanisms underlying both explicit and implicit timing on the scale of seconds continue to be debated. Interestingly however, both fields have independently converged on potentially similar neural mechanisms and signatures, including ramping activity [10–12, 16, 23–26], neural sequences [6, 9, 27–29], and short-term synaptic plasticity [30–34].

Both working memory and timing are functions that operate on the scale of seconds and require information to be transiently stored. During the delay period of standard WM tasks, it is necessary to maintain retrospective information that will be prospectively used. Timing requires tracking elapsed time since a stimulus was delivered (retrospective information) and prospectively estimating when an external event will occur or an internal event should be generated. Recent computational research has suggested that WM and timing could be implemented by the same circuit mechanisms [7, 15, 20, 35]. The shared properties between WM and timing, along with the convergence of potential mechanisms, has led to the suggestion that they may share similar mechanisms, or even that in some cases, they may comprise the same computation [20, 35]. However, *in-vivo* evidence of a shared mechanism for both working memory and timing has yet to be demonstrated in a behavioral task with both WM and timing components.

In animal studies, WM and timing have generally been studied independently of each other and relied on distinct behavioral tasks. To make this link, it is necessary to use a WM task that allows researchers to probe if subjects are learning the temporal structure of the task itself. While most timing research has focused on explicit timing tasks where subjects must keep track of time in order to perform the task, implicit timing tasks provide an opportunity to determine if animals learn the temporal structure of the task, even if the structure is task-irrelevant. Human studies have demonstrated that participants do, indeed, learn the task-irrelevant temporal structure of WM tasks [22, 35–37]. Therefore, we developed a novel olfactory WM and implicit timing task for rodents based on an olfactory Delayed-Non-Match-to-Sample (DNMS) task, in which the identity of the 1^st^ cue signals the duration of the delay period. We refer to this task as a differential-Delayed-Non-Match-to-Sample (dDNMS) task [35]. This task allowed us to demonstrate WM performance deficits when implicit timing expectations were violated, replicating human behavioral results using a similar dDNMS task [35]. We have previously found that during performance of a standard DNMS task, CA1 pyramidal neurons fire sequentially across a fixed 5-second delay following specific odors [9]. We hypothesized that these sequences would be modulated by temporal expectation in our dDNMS task where each odor is followed by a delay period of a distinct duration (2.5 or 5.0 seconds). We found that CA1 odor-specific sequential activity was differently shaped when animals were expecting different delay lengths with an over-representation of CA1 neuronal activity prior to the expected arrival of the 2^nd^ odor, suggesting that implicit timing is encoded within these sequences. Neural trajectories sped up in anticipation of the 2^nd^ odor following a short delay, and this corresponded with better encoding of odor and time on short delay trials. Altogether, we demonstrate that sequential activity in CA1 is a potential shared mechanism for the encoding of working memory and implicit timing.

## RESULTS

We developed a novel working memory and implicit timing task for rodents, by modifying a standard olfactory DNMS working memory task [9, 38]. In this dDNMS task, either odor *A* or odor *B* is presented for 1 second followed by a delay period whose duration depends on the identity of the 1^st^ odor. If the 1^st^ odor presented is odor *A*, the reward period is 2.5 seconds; and if *B* is the 1^st^ odor, the delay is 5 seconds. After the delay period, either odor *A* or *B* is presented as the 2^nd^ odor, and animals are rewarded by responding to the non-match trials following the onset of the 2^nd^ odor. No reward or punishment is provided for match trials. Adult male and female mice (n = 23) were water-restricted and trained to perform the dDNMS task (**Figure 1 A**). Mice were head-fixed above a Styrofoam spherical ball to allow running. After the animals learned the task to criterion (see Methods), we introduced ‘reverse’ trials in which the cue-delay contingency was reversed for 20% of all trials (**Figure 1 A-B**) to investigate whether mice implicitly learned the temporal structure of the task and whether unexpected delay lengths would alter WM performance. Animals performed the task with 80% standard and 20% reverse trials for 2 days, and each day consisted of 7 blocks of 20 trials with pseudorandomly distributed odor combinations.

**Figure 1:**
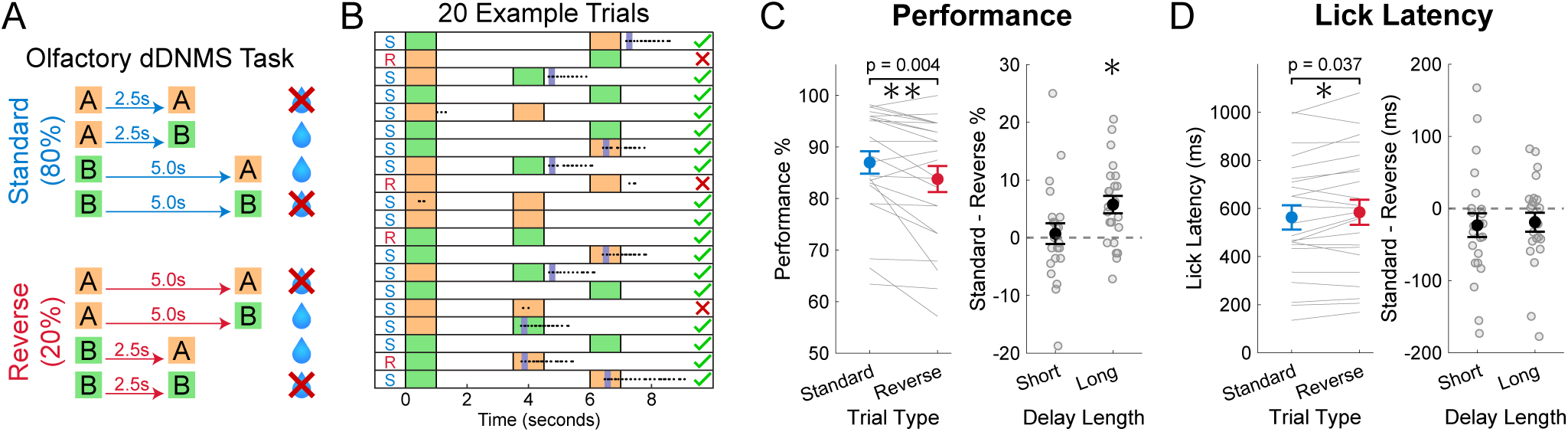
Novel rodent dDNMS working memory task showed behavioral evidence of implicit timing. **A)** Schematic of the olfactory differentially-Delayed-Non-Match-to-Sample (DNMS) task. Water delivery and licking behavior were assessed during a 3-second reward period starting at 2^nd^ odor onset. **B)** Example block of perfect performance for 20 standard (S) and reverse (R) trials. Dots indicate licks and dark blue bars indicate water delivery. **C)** Behavioral performance accuracy calculated as percent-age of trials correct. Large dots represent the mean, and error bars indicate standard error of mean (SEM). All statistics are paired t-tests. The left panel data was split by trial type. The right panel was split by delay length, and the difference between standard and reverse trials is displayed. The asterisk above long delay in right panel indicates significance (p = 0.0009). **D)** Same as (C), but for median lick latency after onset of the 2^nd^ odor.

### Novel rodent dDNMS working memory task showed behavioral evidence of implicit timing

To investigate whether mice showed behavioral evidence of learning the implicit temporal structure of the dDNMS task, we compared performance accuracy and lick latency between standard and reverse trials on the first 2 days of the inclusion of reverse trials. Mice showed decreased performance accuracy on reverse trials compared to standard trials (87.00 ± 10.42 % on standard trials and 83.77 12.10 % on reverse trials, p = 0.0039, **Figure 1 C**). However, this effect was driven entirely by reverse long trials—i.e., trials in which there was a long delay after the ‘short odor’ (86.49 ± 9.47 % on standard long trials and 80.75 ± 14±19 % on reverse long delay trials, p = 0.0009). There was no effect on performance accuracy during the reverse short trials (87.50 ± 11.88 % on standard short trials and 86.80 ± 12.25 % on reverse short delay trials, p = 0.70). Therefore, performance accuracy was only impaired when mice expected a short delay and had to maintain the memory of the 1^st^ odor for longer than anticipated. A potential interpretation of this asymmetry is that in this task, mice learn how long WM should be stored for, and when delays are unexpectantly long, the WM trace is degraded.

Additionally, by measuring the lick latency on each trial when mice made a choice to lick, we found that mice took a longer time to respond on reverse trials (563 ± 243 milliseconds on standard trials and 584 252 milliseconds on reverse trials, p = 0.037, **Figure 1 D**). Interestingly, this effect was driven by both the reverse short and reverse long trials.

Together, the results demonstrate that mice implicitly learned the temporal structure of the differential delays even though the delays are task-irrelevant. Specifically, violating their temporal expectations with reverse trials resulted in both changes in accuracy and lick latency.

### Sequential activity in CA1 reflects implicit learning of the cue-specific differential delay durations

To determine if there was an interaction between the neural signature of WM and implicit timing representations, we performed *in-vivo* two-photon calcium imaging of the dorsal CA1 region of the hippocampus as mice performed the dDNMS task (**Figure 2 A-B**). In our previous work using a standard olfactory DNMS task with fixed 5-second delays, we found that a population of dorsal CA1 neurons fired during specific epochs of the task [9]. While some cells fired during the presentation of specific odors, others fired during specific timepoints of the delay period after a specific odor yielding odor-specific sequential activity tiling the entire delay period [9]. Here, we recorded dorsal CA1 neural activity to investigate how differential temporal expectations influence hippocampal sequential activity.

**Figure 2:**
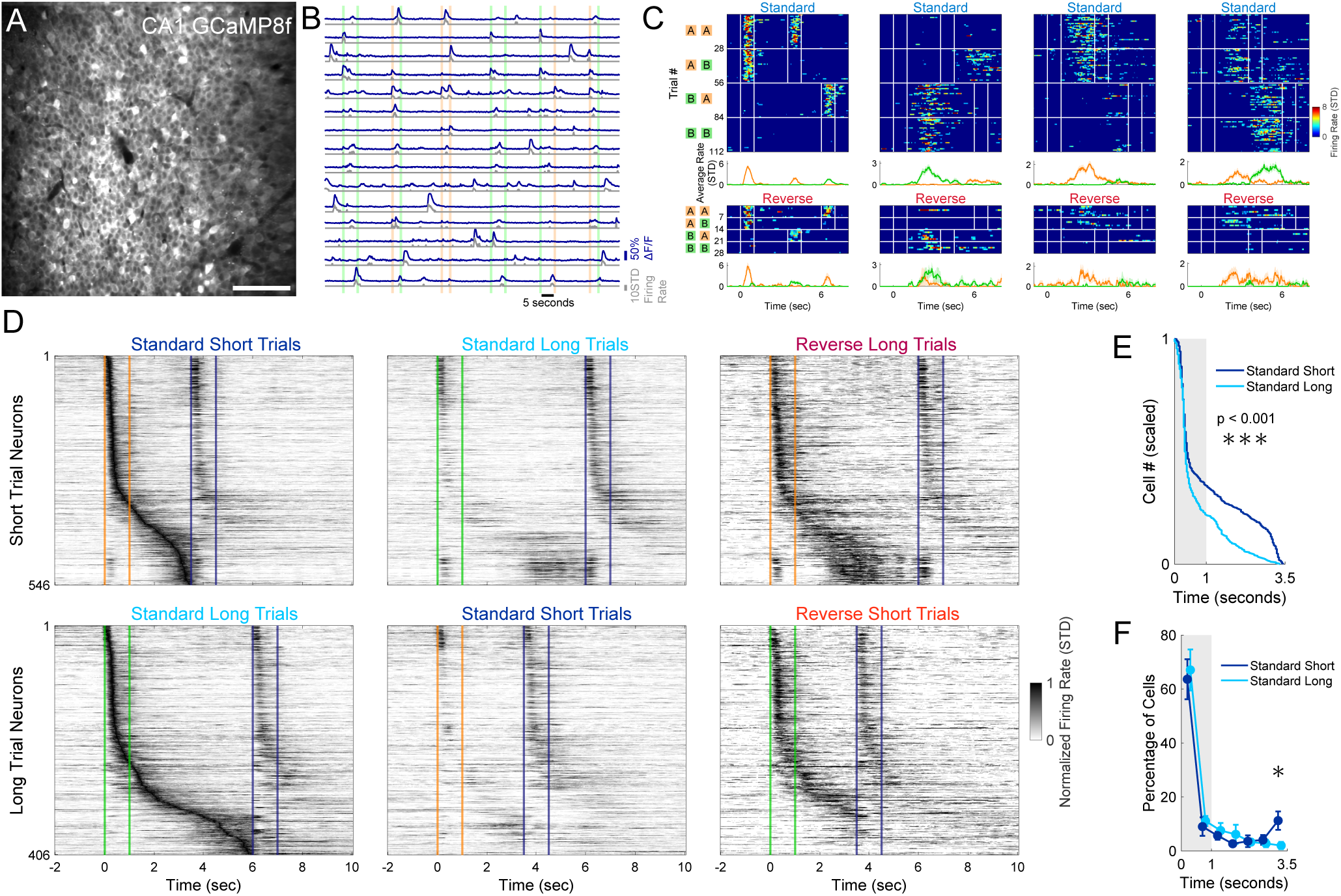
Sequential activity in CA1 reflects implicit learning of the cue-specific differential delay durations. **A)** Example field of view of dorsal CA1 neurons expressing GCaMP8f. Scale bar is 100µm. **B)** Example activity traces of 15 neurons across 6 standard trials. Dark blue traces indicate the ΔF/F, while the gray trace represents the normalized and floored deconvolved signal used to estimate firing rate. Vertical colored bars represent the odor delivery periods. **C)** Heatmaps of four example neurons from the first day including reverse trials. Heatmaps show deconvolved signals on each trial that were grouped into trial type. Vertical white lines indicate onset and offset of each odor delivery. Average traces at bottom display the average firing rate split by trials that started with odor *A* and those that started with odor *B*. **D)** Sequential activity of the neurons that had a significant peak in activity during the 1^st^ odor presentation or delay period. Each row is the average trace normalized and sorted based on its peak firing rate on the preferred trials (left most panels). The top three panels show neurons that have a significant peak during odor *A* or the delay period following odor *A* (short-trial neurons) under the indicated trial types (the middle and right panels are sorted as in the left panel). The bottom three panels are the same but for neurons with a significant peak during or after odor *B* (long-trial neurons). Orange and green lines indicate the 1^st^ odor onset and offset, while blue lines indicate 2^nd^ odor onset and offset (which could be either odor). **E)** Distribution of significant peaks were different between short-trial and long-trial neurons (pooling all cells, Kolmogorov-Smirnov test, p = 4×10^−6^). **F)** Location of significant peaks into 0.5 second bins (n = 11, repeated paired t-tests corrected for multiple comparisons with Benjamini-Hochberg procedure, asterisk indicates p = 0.0315).

Adult male and female mice (n=11) were injected with AAV1-Syn-jGCaMP8f or jGCaMP7f into the right dorsal CA1 and implanted with a 3mm diameter glass-bottomed titanium cannula above the intact alveus after aspiration of the overlying cortex and corpus callosum [9]. Neural activity was recorded for each mouse on its last two days of receiving only standard trials and the first two days with the inclusion of reverse trials, yielding 44 recording sessions with an average of 405 ± 187 active neurons per day (**Figure 2 A-B**). A proportion of CA1 neurons responded reliably during different epochs of the trial. Some had responses with significant activity peaks during the odor presentation, while others had peaks during the delay period (**Figure 2 C**). Altogether, the population of CA1 neurons exhibited sequential activity that tiled the 1^st^ odor presentation and delay period. One group of neurons with significant activity peaks during or after odor *A* formed the short-trial sequence, while another group with peaks during or after odor *B* formed the long-trial sequence. Out of all active neurons, 13.9 ± 12.1 % had significant peaks and were assigned to the short-trial (odor *A*) or long-trial (odor *B*) sequences.

To visualize the sequential activity, we pooled all neurons with significant peaks during the first two days of reverse trials. After grouping short-trial neurons and long-trial neurons separately and then sorting them based on the peak firing, we observed that cells tiled the entire delay period (left panels of **Figure 2 D**). On reverse trials with short delays, the sequence elicited by the long cue was cut short by presentation of the 2^nd^ odor. During reverse trials with long delays—in which the 2^nd^ odor exceeded the expected delay—the neural sequence elicited by the short cue appeared to end prematurely failing to reach the onset of the 2^nd^ odor (right panels of **Figure 2 D**).

Visual inspection of **Figure 2 D** suggests an over-representation of the end of the short delay, and a modest over-representation of the end of the long delay. To quantify if differences in expected delay duration influenced the structure of sequential activity, we compared the standard short-trial and standard long-trial neural trajectories during the first 3.5 seconds of the trials (encompassing the 1^st^ odor presentation and the short delay length). This allowed a comparison of equivalent time lengths to determine if the hippocampus allocates additional resources to the anticipation of the 2^nd^ odor arrival in the short trial sequence. The distribution of these pooled significant neurons was different between the short and long trial sequences (Kolmogorov-Smirnov test p = 4×10^−6^, **Figure 2 E**). This effect was driven by the large over-representation in the last 0.5 seconds of the short delay period (3.0 to 3.5 seconds trial time) for the short trial sequence neurons, with 11.2 ± 11.4 % of short trial neurons with a peak during the last 0.5 seconds compared to only 2.0 ± 4.5 % for long trial sequence neurons (out of only neurons with a peak during the first 3.5 second period, n = 11, paired t-test p = 0.0315, **Figure 2 F**). Together, these findings suggest that odor-specific sequential activity in CA1 reflected both the WM of the 1^st^ odor identity and the temporal expectation of the 2^nd^ odor.

### Neural trajectories increase in speed in anticipation of the 2^nd^ odor

To further quantify the differences between CA1 population activity elicited by the cues associated with short and long delays, we evaluated neural trajectory dynamics of all significant short-trial or long-trial sequence neurons. First, we performed principal component analysis (PCA) and visualized the first two principal components (PCs) of population activity during the 1^st^ odor and delay period. By splitting the neural trajectories based on the trial type, we revealed differences in neural trajectory speed and distance from a baseline resting state before the presentation of the 1^st^ odor (**Figure 3 A**). Following rapid changes in population activity during the odor presentation, neural trajectories slowed and returned toward the baseline state. However, on standard short trials, the trajectory turned away from the baseline and increased its speed as if in anticipation of the 2^nd^ odor (**Figure 3 B and D**). When comparing the trajectories of standard short and reverse short trials, the difference was significant during the last second of the short delay. On reverse long trials, when the 2^nd^ odor comes later than expected, the trajectory slowed and returned toward baseline (**Figure 3 A, C, and E**). When comparing standard long and reverse long trials (both with 5 second delays), the distance from baseline and trajectory speed was significantly higher for reverse long trials in the middle of the delay period when the 2^nd^ odor was expected. This is likely driven by the over-representation of significant sequence cells that have their peak activity at the end of the short delay following odor *A*. Collectively, these findings corroborate that CA1 neural trajectories have differences that reflect the learned cue-delay associations.

**Figure 3:**
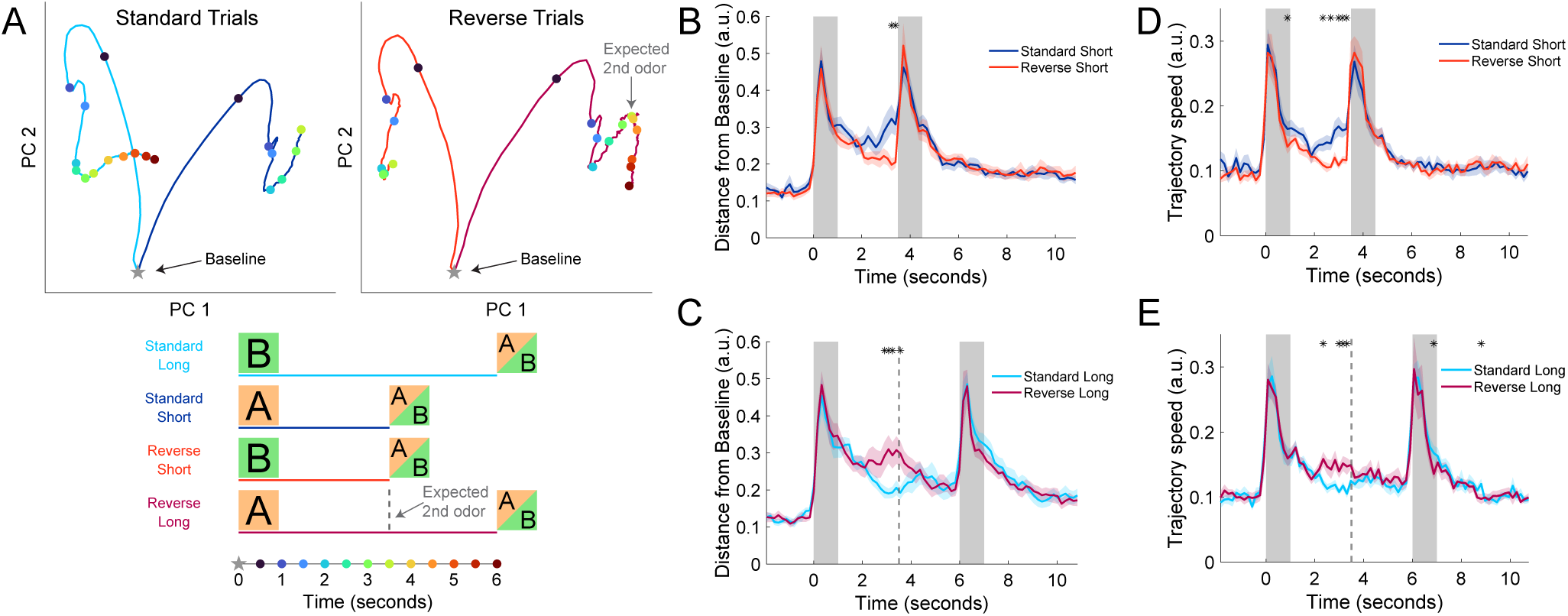
Neural trajectories increase in speed in anticipation of the 2^nd^ odor. **A)** Principal component analysis of neural trajectories during 1^st^ odor and delay period. Top left panel are the averaged trajectories of standard long (cyan) and standard short (blue) trials, while top right panel are reverse short (red) and reverse long (magenta) trials. Bottom shows time points of colored dots that correspond to half-second increments in top panels. Baseline was calculated as the activity that precedes the 1^st^ odor (from -2 to 0 seconds). Dotted line for ‘expected 2^nd^ odor’ refers to the expected arrival of 2^nd^ odor of reversal long trials following the cued-short odor *A*. **B)** Distance from baseline was calculated as the Euclidean distance between neural activities at a point in time compared to a 2-second baseline period before the 1^st^ odor. In the comparison of standard short and reverse short trials, since there are many more standard than reverse trials, the analysis was balanced by taking the standard trials nearest to each reverse trial. The population of significant short-trial neurons (from Figure 2 D-F) was used for the standard short trace, and the significant long-trial neurons were used for the reverse short trace. Traces from each recording were normalized by the number of significant neurons (n) by dividing traces by √n. Thick lines represent the mean of 22 recording sessions (first two days with inclusion of reverse trials), and shaded area represents standard error of the mean. Asterisks represent bins of 1/6 second that were significantly different (Two-Way ANOVA animal and day, corrected for multiple comparisons with Benjamini-Hochberg procedure, p < 0.01). **C)** Same as (B) but comparing the standard and reverse trials with long delays. The dotted line is the same as (A) for expected 2^nd^ odor arrival on reverse long trials. **D-E)** Same as (B-C) but for trajectory speed, which is the Euclidean distance between neighboring bins.

### Running patterns showed evidence of implicit timing but did not explain effects of neural trajectories

Since mice could locomote on the spherical Styrofoam treadmill during trials, we next asked whether locomotion also exhibited evidence of implicit timing and whether neural trajectories reflected locomotion patterns rather than timing. While mice exhibited a range of movement patterns with some rarely moving on the treadmill, most mice primarily flinched or twitched at the onset of odor presentation and only occasionally ran during the delay and reward periods (**Supplemental Figure 1**). When comparing standard short and standard long trials, mice showed an anticipatory increase in their overall locomotion and running earlier in standard short trials (**Supplemental Figure 1 A**). This difference was statistically significant for all bins of the short delay period as mice increased their locomotion immediately following the 1^st^ odor only on standard short trials. Comparing standard short and reverse short trials showed the same effect; however, increased variation due to fewer trials reduced our power to detect significant differences as early in the delay period (**Supplemental Figure 1 B**). Most strikingly, mice continued to increase their locomotion on reverse long trials until the late arrival of the 2^nd^ odor (**Supplemental Figure 1 C**). This late period following the missed expected 2^nd^ odor was the period with the greatest locomotion and running of all trial types and time periods.

Despite running occurring only in a minority of trials, the behavioral differences between delay periods led us to evaluate if these differences could explain the effects seen in the neural trajectories. Firstly, as shown in **Figure 3 B and D**, in standard short trials, distance from baseline and trajectory speeds decreased early in the delay and only increased and became statistically significantly greater than standard long trials during the last second of the short delay. This is different from locomotion and running which steadily increased during the entire short delay period (**Supplemental Figure 1 A and B**). Secondly, as shown in **Figure 3 C and E**, in reverse long trials, distance from baseline and trajectory speeds were only increased above standard-long trials in the middle of the delay at the expected 2^nd^ odor arrival. Conversely, the locomotion and running during reverse long trials continued to increase until the eventual arrival of the 2^nd^ odor (**Supplemental Figure 1 C**). As a final control to ensure that running did not drive neural trajectory effects, we analyzed trajectory speed and distance from baseline in standard short and standard long trials when only non-running trials were included. The significant increase in speed and distance were found similarly with and without the inclusion of running trials (**Supplemental Figure 2**). Together these results provide further evidence that mice learned the implicit temporal structure of the task by increased locomotion in anticipation of the 2^nd^ odor, but locomotion cannot account for the differential neural trajectories observed in the delay periods.

### CA1 encoding of odor presentation and elapsed time is stronger in response to the short delay cue

Finally, we asked if the different neural trajectories, elicited by odors preceding short and long delays, yield a difference in the strength of odor encoding and encoding of elapsed time.

To answer the question of encoding odor, we calculated mutual information in bits per cell about whether an odor had been presented or not. Specifically, we estimated the amount of information between a baseline bin (1.5 seconds before 1^st^ odor) to all other time bins. This measure of information was averaged for each neuron to estimate the strength of the encoding of an odor presentation. As expected, during the first two days with reverse trials, odor information was greatest during odor presentation periods; importantly however, odor information increased at the end of the short delay of standard trials in anticipation of the 2^nd^ odor (**Figure 4 A**). The same effect was seen in the middle of reverse long trials following the cued-short odor when odor information increased significantly higher than that of standard long trials (**Figure 4 B**).

**Figure 4:**
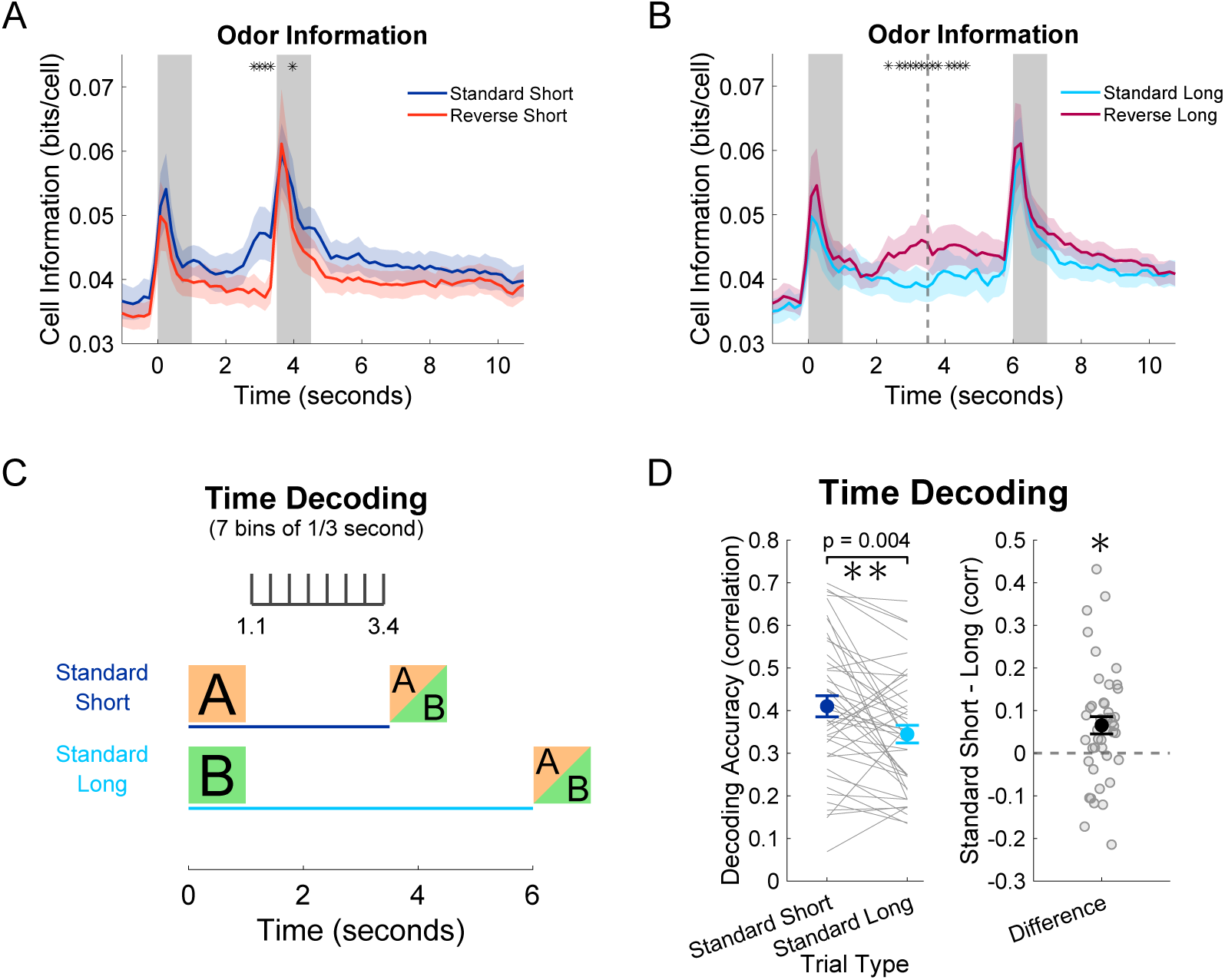
CA1 encoding of odor presentation and elapsed time is stronger in response to the short delay cue. **A)** Odor information comparing baseline bin (-1.5 seconds) to all other bins in standard and reverse trials with short delays. Since there are many more standard than reverse trials, the analysis was balanced by taking the standard trials nearest to each reverse trial. Thick lines represent the mean of 22 recording sessions (first two days with inclusion of reverse trials), and shaded area represents standard error of the mean. Asterisks represent bins of 1/6 second that were significantly different (Two-Way ANOVA animal and day, corrected for multiple comparisons with Benjamini-Hochberg procedure, p < 0.01). **B)** Same as (A) but for long delay trial types. The dotted line represents the expected 2^nd^ odor arrival on reverse-long trials. **C)** Multiclass support vector machine (SVM) decoding of time bins within the short delay period between 1.1 and 3.4 seconds with 7 bins each lasting 0.33 seconds. **D)** Decoding accuracy calculated as the correlation of real bins to the decoder predicted bins (0 being chance, and 1 being perfect). Two-Way ANOVA animal and day for the 44 recording sessions, p = 0.004).

To answer the question of decoding elapsed time, we trained a support vector machine (SVM) to decode time bins based on neural population activity. To allow a fair comparison between short and long trials, we only performed decoding during the short delay window from the 1.1 to 3.4 seconds in order to exclude any directly cue-related activity (**Figure 4 C**). Decoder performance was quantified as the correlation of the actual and predicted bins based on the neural trajectories of standard short and standard long trials. Across all four days of recording (last two days of only standard trials and first two days with reverse trials), time decoding was higher in the standard short trials (**Figure 4 D**).

Together, these findings indicate that in the first 2.5 seconds of the delay periods, CA1 population activity more strongly encodes odor and time when expecting a short delay. This improvement is likely to be driven in part by the need to anticipate and prepare for the onset of the 2^nd^ odor. These results further establish that in addition to the differential structure of the neural trajectories elicited by the short and long delay cues, there are functional differences in how well these trajectories encode odor and time.

## DISCUSSION

Using a novel rodent olfactory-based working memory (WM) task, we demonstrate that mice implicitly learned the task irrelevant cue-delay associations as revealed by a decrease in accuracy when delay length expectations were violated. These results are generally consistent with previous WM and implicit timing experiments in humans [22, 35, 39] and comprise one of the first examples of implicit timing in a rodent task. Using two-photon calcium imaging of dorsal CA1 pyramidal neurons during performance of the dDNMS task, we found odor-specific sequential activity encoding of WM and elapsed-time representations during the delay periods. First, odor-specific sequential activity tiled the entire delay period, allowing the 1^st^ odor representation to be maintained until the arrival of the 2^nd^ odor. Second, sequential activity also encoded odor-specific elapsed time as different neurons became active at different time points within the delay period. Third, the speeds and shapes of the neural sequences were different for the short and long delay cues, establishing that the temporal structure of the tasks differentially sculpted the neural trajectories generated by each cue. Finally, during the short delays, there was an over-representation of activity in windows prior to the arrival of the 2^nd^ odor, which corresponded with an improvement in the ability to decode both odor and elapsed time.

### Does WM expire?

Our behavioral results revealed a decrement in accuracy during unexpected delays. Importantly, this effect was not symmetric for accuracy. Compared to standard trials, there was no decrease in performance during reversal trials in which the cue indicated a long delay but a short delay occurred. In contrast, performance was impaired when a short delay was expected, but a long delay was presented. The cause of the observed asymmetry is unclear. If the performance decrement was entirely due to implicit timing—e.g., not being prepared to access information in WM or to process the 2^nd^ odor [22]—the reversal effect should occur for both unexpected short and long delays. Indeed, the reaction time effect was symmetric, suggesting that implicit timing contributes to anticipatory attention or motor preparation [18, 40].

One interpretation of the observed asymmetry for accuracy is that the duration of encoding of WM itself is tuned in a task dependent manner. For example, when information must be stored in WM for a known period of time, WM could “expire” in a learned fashion to optimize cognitive resources—i.e., WM for the short-cue could be shorter lasting than for the long-cue, because animals know a priori for how long the memory will have to be stored. While studies have shown that WM can decrease at longer intervals [41, 42], to the best of our knowledge, few studies have directly addressed the duration of WM in tasks which subjects first learn the expected delay duration and then are challenged with new durations, as in the dDNMS task. This approach, however, is constrained by the fact that subjects will quickly learn the unexpected delays—a factor that could have limited our ability to detect larger effects.

The notion that animals may learn to tune WM to expire in a task dependent manner has profound implications for theories of WM and explains some of the diversity of the dynamic regimes observed during WM tasks. For example, an ongoing controversy pertains to the neural dynamic regimes used to store WM, including persistent stationary activity, dynamic neural trajectories (e.g., ramping and neural sequences), or activity silent mechanisms [32, 43, 44]. Time-varying regimes such as neural sequences and ramping activity provide a clear mechanism for the learned, timed expiration of WM as they have natural end or saturation points. Thus, it may be the case that differences in the observed dynamic regimes associated with WM reflect, in part, differences in the temporal structures of the task. Indeed, rodent studies have reported different neural regimes during the delay period of tasks depending on whether a fixed or randomized delay was used [8, 14].

### Neural sequences in WM and timing tasks

Neural sequences have been reported in both WM [8, 9, 45, 46] and timing tasks [6, 47–50]. To date, however, studies have not used tasks that allow for dissociation between the WM and temporal components of behavioral tasks. Here, by using the dDNMS task we were able to establish that neural sequences are shaped by task structure, including an increase in trajectory speed at the expected time of the 2^nd^ odor in the standard short delay trials. The differential dynamics of the neural sequences resulted in improved decoding of time in response to the short cue during the 1.0-3.5 s delay period (i.e., during the time window shared between both delays).

It has been proposed that neural sequences provide an ideal neural dynamic regime to encode time-varying information because their relatively high dimensionality and orthogonality provides a flexible way for downstream neurons to readout information using biologically plausible learning rules [49, 51]. Neural sequences have been observed in numerous brain areas including the striatum, neocortex, hippocampus, cerebellum, and area HVC in zebra finches [6, 9, 27, 28, 47, 48, 52–55]. Notably, many of these areas exhibit highly distinct microcircuit architecture. It is thus likely that while some areas are intrinsically capable of generating neural sequences, other areas are reading out the dynamics generated upstream or simply relaying these neural trajectories. Dorsal CA1 was chosen for this study for its involvement in WM and representations of time through sequential activity [6, 9, 56–58]; however, its relatively weak recurrent connectivity [59–61] makes it unlikely to be generating the temporal representations, as computational models that generate multiple neural sequences generally rely on the recurrent excitation present in neocortical circuit [7, 35, 62–64].

The odor-specific working memory information observed in CA1 is likely integrated from multiple areas such as lateral entorhinal cortex [65–68] and piriform cortex [69–71], but it is unclear which brain regions play a role in maintaining this information during the delay period. The CA3 region of the hippocampus—given its stronger recurrent connectivity and inputs to CA1 [72, 73]—may play a critical role in the formation of the sequential activity we observed in CA1. Alternatively, both the medial and lateral entorhinal cortices have been implicated in timing representations as well [54, 74, 75]. Whatever the mechanisms generating these neural sequences, the current study does make it clear that they are not hardwired dynamic regimes, but are shaped by the task at hand, as revealed by the differential speed and shape of the neural trajectories evoked by the short- and long-cues. Future manipulation studies will be necessary to understand which regions play a role in generating the timing-dependent dynamics we have recorded in this study.

## RESOURCE AVAILABILITY

### Lead Contact

Further information and requests for resources and reagents should be directed to the lead contacts, Dean Buonomano and Peyman Golshani.

### Material Availability

No new materials were created for this study.

### Data and Code Availability

Data and analysis code will be made available on Dryad and GitHub after manuscript acceptance. Any additional information necessary to reanalyze the data reported in this paper is available from the lead contacts.

## ACKNOWLEDGMENTS

We thank K. Reyes, B. Madruga, Z. Day, and X. Chen for technical support. This work was supported by the following funding sources: T32DGE1829071, 5T32NS045540-20, R01NS116589, and 1P50HD103557-01.

## AUTHOR CONTRIBUTIONS

C.D., J.T., D.B., and P.G. conceived the experiments. C.D. conducted experiments. C.D. and D.B. analyzed experimental data. C.D. generated figures. C.D., D.B., and P.G. wrote the manuscript.

## EXPERIMENTAL MODEL AND SUBJECT DETAILS

### Animals

All of the experiments were conducted according to the National Institute of Health (NIH) guidelines and with the approval of the Chancellor’s Animal Research Committee of the University of California, Los Angeles. A total of 9 adult male and 3 female mice (8-34 weeks old) were used for *in-vivo* behavioral experiments, and a total of 6 adult male and 5 female mice (8-16 weeks old) were used *in-vivo* calcium CA1 neuron imaging experiments. All were C57BL/6J (Jackson Laboratory, 000664), experimentally naïve, and housed in the vivarium under a 12-hour light/dark cycle. All mice were group housed (2-4 per cage) with the exception of 3 that had to be separated following surgery because of fighting.

## METHOD DETAILS

### Surgical Procedures

Mice (8-30 weeks old) were subcutaneously administered pre-operative drugs (carprofen 5 mg/kg, dexamethasone 0.2 mg/kg, lidocaine 5 mg/kg) 30 minutes before surgery. Mice were anaesthetized with isoflurane (5% induction, 1-2% for maintenance), and anesthesia was continuously monitored and adjusted as necessary. The scalp was shaved, and mice were placed into a stereotactic frame (David Kopf Instruments, Tujunga, CA) on a feedback-controlled heating pad (Harvard Apparatus) set to maintain body temperature at 37*^◦^*C. Eyes were protected from desiccation using artificial tear ointment. The surgical incision site was cleaned three times with 10% povidone-iodine and 70% ethanol. Fascia was removed by applying hydrogen peroxide, connective tissue was cleared from the skull, and the skull was scored to facilitate effective bonding with adhesives at the end of surgery.

Behavioral experiments: After removing fascia and connective tissue, a custom-made lightweight stainless-steel headbar was attached to the posterior skull and secured with cyanoacrylate glue. Dental cement (Ortho-Jet, Lang Dental) was applied to seal and cover any remaining skull.

CA1 calcium imaging experiments: After stereotactically aligning the skull, a single burr hole was made above right dorsal CA1. Either pGP-AAV1-syn-jGCaMP8f-WPRE (1000nL of 1:5 saline dilution) or pGP-AAV1-syn-jGCaMP8f-WPRE (1000nL of 1:5 saline dilution) was injected into the right dorsal CA1 (2.0 mm posterior from bregma, 1.8 lateral from bregma, and 1.3 ventral from dura). Viruses were injected using a Nanoject II microinjector (Drummond Scientific) at 60nL per minute. Following virus injection and waiting for 45 minutes, a circular craniotomy (3 mm diameter) was made centered around a point made 2.0 mm posterior and 1.8 lateral to bregma. Dura beneath the craniotomy was removed and cortical tissue above dorsal CA1 was carefully aspirated using a 27-gauge blunt needle. Corpus callosum was spread to the sides of the craniotomy to expose the alveus. Cortex buffer (NaCl = 7.88g/L, KCl = 0.372g/L, HEPES = 1.192g/L, CaCl_2_ = 0.264g/L, MgCl_2_ = 0.204g/L, at a pH of 7.4) was continuously flushed during aspiration and until bleeding stopped. A titanium ring with a 3 mm diameter circular thin #0 coverglass attached to its bottom was implanted into the aspirated craniotomy and the overhanging flange was secured to the skull with vetbond (3M). A custom-made lightweight stainless-steel headbar was attached to the posterior skull and secured with cyanoacrylate glue. Dental cement (Ortho-Jet, Lang Dental) was applied to seal and cover any remaining skull, and to form a small well around the titanium ring for holding immersion water for the objective during imaging.

Following surgery, all animals were given post-operative care (carprofen 5 mg/kg and dexamethasone 0.2 mg/kg for 48 hours after surgery) and provided amoxicillin-treated water at 0.5 mg/mL for 7 days. All mice recovered for 7-14 days before experiments began.

### Experimental setup

The entire behavioral setup is as described in Taxidis et al. [9]. Mice were head-fixed above an 8-inch spherical Styrofoam ball (Graham Sweet) which can rotate about one axis for 1D locomotion that was recorded with a sensor (Avago ADNS-9500). A continuous stream of clean air (~1 L/min) was delivered toward the animal’s nose via Tygon PVC clear tubing and a custom-made port that held the air tube and water port. At the onset of the odor presentation period, a dual synchronous 3-way valve (NResearch) switched to the odorized one for 1 second. Odorized air was created by using a 4-port olfactometer (Rev. 7c; Biology Electronics, Caltech) supplying air to either of two glass vials containing odor A (70% Citral, FCC; Sigma Aldrich) or odor B ((-)-a-Pinene ± 97%, FCC; Sigma Aldrich), which were both diluted in mineral oil at 5% concentration. Water droplets (~10µl) were released by a 3-way solenoid valve (Lee Company), and licks were detected by using a custom battery-operated circuit board with one end of the circuit connected to the headbar and the other to the lickport. The behavioral rig was controlled with custom written software (MATLAB) and through a data acquisition board (USB-6341: National Instruments).

### Behavioral training

After 7-14 days recovering from surgery, mice were handled and began water-restriction to 85% of their original weight before water-restriction. After one day of handling, mice were habituated to being head-fixed above the spherical treadmill (can rotate about one metal axis for 1D locomotion that is recorded) for two days. On the 4th day of training, mice began learning to lick from the lickport as water was automatically delivered at the beginning of the reward period following only standard non-matched odor trials (AB or BA, with water delivery at the offset of the 2^nd^ odor). Trials were delivered in blocks of 20 trials. This phase was always 2 days except for the rare mouse that needed one extra day to reach motivation level and lick water from the port for at least 50 trials. In the next phase, water was only delivered if the mouse licked during the response period, and mice learned to reliably lick in anticipation of the reward following the 2^nd^ odor. This phase was also 2 or 3 days, dependent on licking during the response period of at least 50 trials. The next phase was the full differential delayed non-match-to-sample (DNMS) task in which standard matched odor trials (AA and BB) were introduced and mice learned to refrain from licking the port following these trials. There was no punishment or timeout following an incorrect lick; the water was simply not delivered. For this phase and previous phases, the response window was 3 seconds starting at the offset of the 2^nd^ odor. After 6 days of this phase or until behavior reached 85 %, we began sliding the response window forward by 1/3 of a second. For example, the first day with the new response window started 2/3 of a second after the onset of the 2^nd^ odor. On the following day, the response window would start 1/3 of a second after the onset of the 2^nd^ odor. Finally, mice performed the dDNMS task with the response window starting at the onset of the 2^nd^ odor. At this stage, 7 blocks of 20 trials were always delivered. Mice performed 3 days of this phase with only standard trials delivered, except for 3 mice that were given an extra 4th day because of performance below 85%. This phase is referred to as the standard trial days. Following this phase, reverse trials were introduced at a 20% rate for 3 more days. Again, 7 blocks of 20 trials were delivered. For simplicity throughout the figures and text, odor A (citral) always predicts short delay and odor B (pinene) always predicts long delay. However, we counterbalanced mice with half trained the opposite way with odor A (citral) predicting long delay and odor B (pinene) predicting short delay.

### *In-vivo* two-photon imaging

All two-photon calcium imaging was conducted using a resonant scanning two-photon microscope (Scientifica) fitted with a 16x 0.80 NA objective (Nikon) to record 512×512 pixel frames at 30.9 Hz. CA1 imaging fields of view were 500×500 µm and axonal imaging fields were 250×250 µm. Excitation light was delivered with a Ti:sapphire excitation laser (Chameleon Ultra II, Coherent), operated at 920 nm. GCaMP8f and GCaMP7f fluorescence was recorded with a green channel gallium arsenide photomultiplier tube (GaAsP PMT; Hamamatsu). Microscope control and image acquisition were performed using LabView-based software (SciScan). Imaging and behavioral data were synchronized by recording TTL pulses generated at the onset of each imaging frame and olfactory stimulation digital signals at 1 kHz, using WinEDR software (Strathclyde Electrophysiology Software).

For imaging experiments, dorsal CA1 was imaged for at least 5 consecutive days of task performance. This includes 2 days of standard trials only and 3 days with reversal trials (however, the last reversal trial day was not included in any analysis presented here). While careful attention was given to aligning the FOV to the previous day’s when possible, sometimes FOV’s needed to be changed to optimize the number of active neurons. We used rotating stages, a motor for adjusting mouse head angle, and a tiltable objective attachment with two degrees of freedom to fine-tune the alignment. Laser power and PMT settings were kept consistent between days, except for rare occasions when it was necessary to keep similar signal-to-noise.

For each day of recording, imaging was halted between each of the 7 blocks of 20 trials. This allowed fine-tuning of alignment to keep the same FOV within the day, and it also prevented brain heating or photo-toxicity. Laser power was kept as minimal as possible (60-90mW) without sacrificing signal-to-noise ratio, and no significant photo-bleaching was observed.

### Histology

Following all experiments, mice were deeply anaesthetized under isoflurane and transcardially perfused with 30 mL 1x PBS followed by 30 mL 4% paraformaldehyde in 1x PBS at a rate of approximately 4 mL/min. After perfusion, the brains were extracted and post-fixed in 4% paraformaldehyde. Coronal sections of 80 µm were collected using a vibratome, 24-48 hours after perfusion. The sections were mounted onto glass slides and cover-slipped with DAPI mounting medium. Images were acquired on an Apotome2 microscope (Zeiss; 5x, 10x, 20x objectives) to confirm proper expression and location of viral expression. For CA1 imaging experiments, GCaMP8f or GCaMP7f was confirmed to be in dorsal CA1 with no damage to the hippocampal formation.

## QUANTIFICATION AND STATISTICAL ANALYSIS

### Performance and lick-latency

For all 23 mice (behavior only and CA1 imaging batches), performance was measured as the percentage of trials with the correct response. Responses were only considered for performance during the response window (3 second period starting at the onset of 2^nd^ odor). Within the response period, the time elapsed between the onset of the 2^nd^ odor and the first lick is considered the ‘lick latency’. To minimize the effect of extremely late licking outlier events, the median lick latency was taken for each recording session.

### Calcium imaging data pre-processing

For CA1 imaging experiments, each day was analyzed separately as neurons were not aligned across days. Movies were processed using the Python implementation of Suite2P 0.9.2 [76] to perform non-rigid motion registration, neuron segmentation, extraction of fluorescence signals, and deconvolution with parameters optimized to our GCaMP8f and GCaMP7f CA1 recordings (with only the decay time-constant being different between the two). We used the default classifier and an ‘iscell’ threshold of 0.3 to only include masks that were likely neurons. Deconvolved signals were taken as the selected output from Suite2P and taken to MATLAB 2021a for further processing. Deconvolved signals were smoothed by a rolling mean of 10 frames (0.32 seconds). To avoid including small events within the range of noise, we performed a hard threshold by z-scoring the deconvolved signal and values below 2 were set to zero [77, 78]. The resulting signal was what was used for all analysis and referred to as ‘firing rate (STD)’ as a proxy for spiking activity [9]. Signals were aligned to the trial structure (odor presentations and lick timing) and the recorded locomotion as mice ran on the spherical ball.

### Sequence neuron detection and analysis

To evaluate peak firing timing, we performed sequence detection similar to the previously described approach in our DNMS task [9]. First, standard trials that begin with Odor A and those that begin with Odor B are separated, and the one with a larger peak of the average activity was considered further. Additionally, only the 3.5 or 6-second period including 1^st^ odor presentation and the delay period was considered. The peak of average activity within this period and a given trial type was determined to be significant if the peak was greater than the 95^th^ percentile of 2000 circular shuffles. The cell must also have had a trial reliability of at least 20% and have fired above 2 STD for 20% of the preferred trials within 0.5 seconds of the peak frame found in the previous step. If a cell passed both criteria, it was considered to be a ‘short trial neuron’ or ‘long trial neuron’ regardless of its odor selectivity.

### PCA analysis and Trajectory Speed

For Figure 3 A, all active neurons were taken, and principal component analysis (PCA) was conducted on the average deconvolved signal of the concatenated 4 different trial types. Trial types were balanced in the number of trials by taking the nearest neighbor standard trial for each reverse trial. The baseline point was calculated as the average position of the trajectories in the 2 seconds preceding the onset of the 1^st^ odor. While Figure 3 A only shows the first 2 principal components (PCs), Figure 3 B-E calculates the Euclidean distances without any dimensionality reduction. These figure panels only use the significant ‘short trial neurons’ and ‘long trial neurons’ detected in the previous figure. Using bins of 5 frames (0.16 seconds), distance from baseline is defined as the distance in full dimensional space between the baseline point and the bin in question. Trajectory speed is the distance in full dimensional space from one bin to the next. Arbitrary units (a.u.) are displayed because distance and speed are normalized for the number of significant neurons (n) by dividing by √n.

### Locomotion analysis

1D locomotion that was recorded with a sensor (Avago ADNS-9500) at 1kHz and binned to match the frame rate of calcium imaging. Signals were binned into 5 frames (0.16 seconds), then z-scored, and finally values below 1 were set to zero. These binned signals are displayed as ‘locomotion (z-score)’. Since most of the locomotion was small movements around the onset and offset of odors, in other analysis we binarized locomotion into ‘not running’ and ‘running’ bouts. A bout of running must have been at least 1 second of locomotion values above 1; and all other periods were considered to be ‘not running’. A trial was considered to be ‘running’ if a 1 second bout of running occurred during the delay period.

### Odor Information

Firing rates were binned into time bins (5 frames, 0.16 seconds) and additionally discretized into 20 activity bins based on the maximal value of each neuron. The mutual information between a baseline bin at 1.5 seconds before the 1^st^ odor was compared to each other time bins between -1 and 11 seconds in trial time separately for each trial type. As in the previous section, trial types were balanced in the number of trials by taking the nearest neighbor standard trial for each reverse trial. For each bin, the resulting values of each neuron were averaged.

### Time decoding

As described in the text, support vector machine (SVM) error-correcting output codes (ECOC) multiclass decoding was used to decode the time bin within the short delay period between 1.1 and 3.4 second time points with 7 bins of size 1/3 seconds. The MATLAB function ‘fitcecoc’ was used to perform the decoding using the default one vs one method. As described previously, the correlation between actual and predicted bins was used to quantify the performance of the decoder [28].

### Statistical analysis

All statistical tests are described in the legend text. For all figures, individual p-values are listed in the figure or legend text. On occasions when single asterisks were displayed above a curve or trace, p-values were corrected for multiple comparisons using the false discovery rate Benjamini-Hochberg procedure, and asterisks indicate bins in which the adjusted p-value < 0.01. In all cases in the text, values were written in the format ‘mean ± standard deviation’ (STD), while error bars in all figures show the mean and standard error of the mean (SEM). No statistical methods were used to determine appropriate sample sizes but were chosen as being comparable to sizes used in similar publications.

## SUPPLEMENTAL FIGURES

**Supplemental Figure 1:**
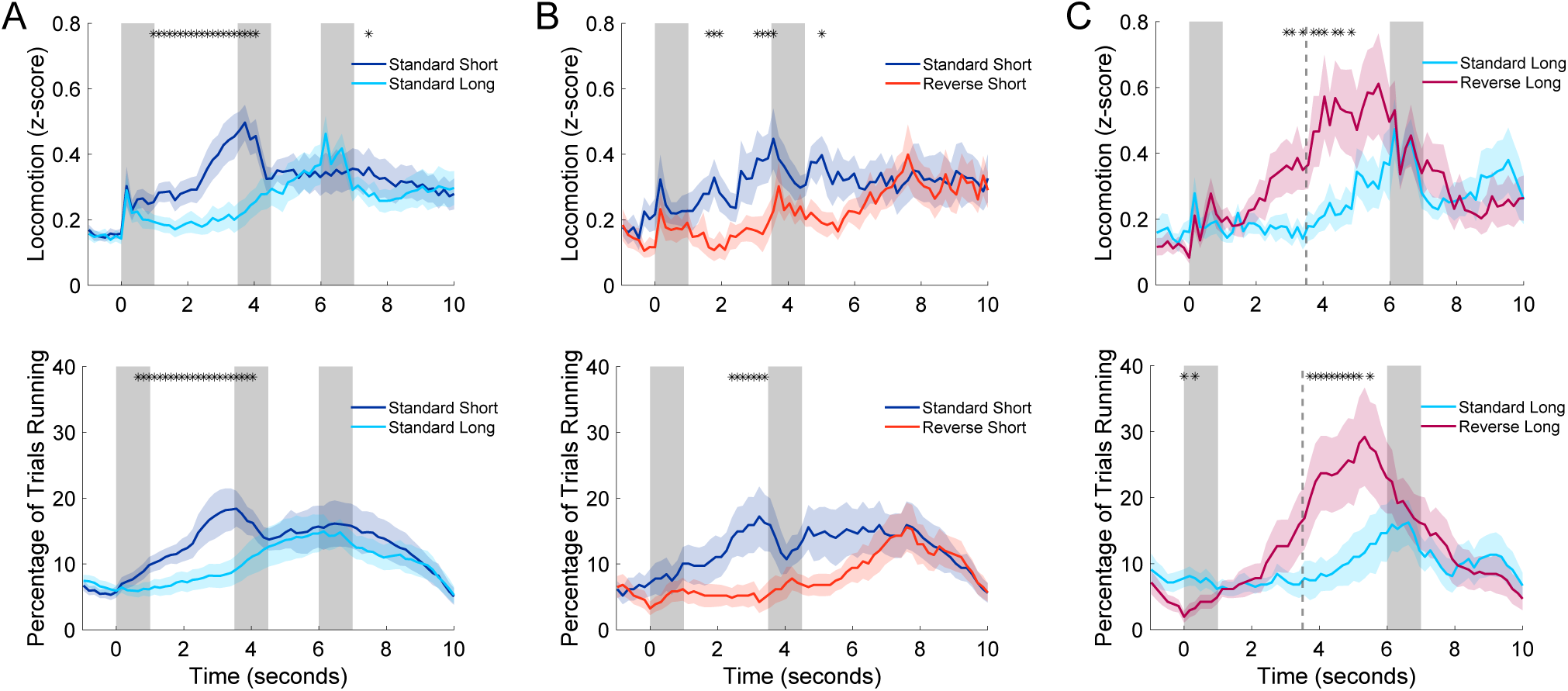
Running differences during delays showed evidence of implicit timing. **A)** Top panel: locomotion is z-scored voltage signal from motion sensor binned to 1/6 second. Bottom panel: percentage of trials with running in a given bin. For both panels, comparison was made between all standard short and standard long trials from the last 2 standard days and first 2 reverse days. The gray bar at time point 0 is the 1^st^ odor for both trial types, while the one at 3.5 is the 2^nd^ odor for standard-short trials and the one at 6 is the 2^nd^ odor for standard-long trials. Thick lines represent the mean of 44 recording sessions (11 mice across 4 days), and shaded area represents standard error of the mean. Asterisks represent bins of 1/6 second that were significantly different (Two-Way ANOVA animal and day, corrected for multiple comparisons with Benjamini-Hochberg procedure, p < 0.01). **B)** Same as (A), but comparison between standard-short and reverse-short trials from first 2 reverse days with the number of trials balanced by taking the nearest neighbor standard trial for each reverse trial. Statistics were the same as (A), but for only 22 recording sessions. **C)** Same as (B), but comparison between standard-long and reverse-long trials. The dotted line represents the expected 2^nd^ odor arrival on reverse-long trials.

**Supplemental Figure 2:**
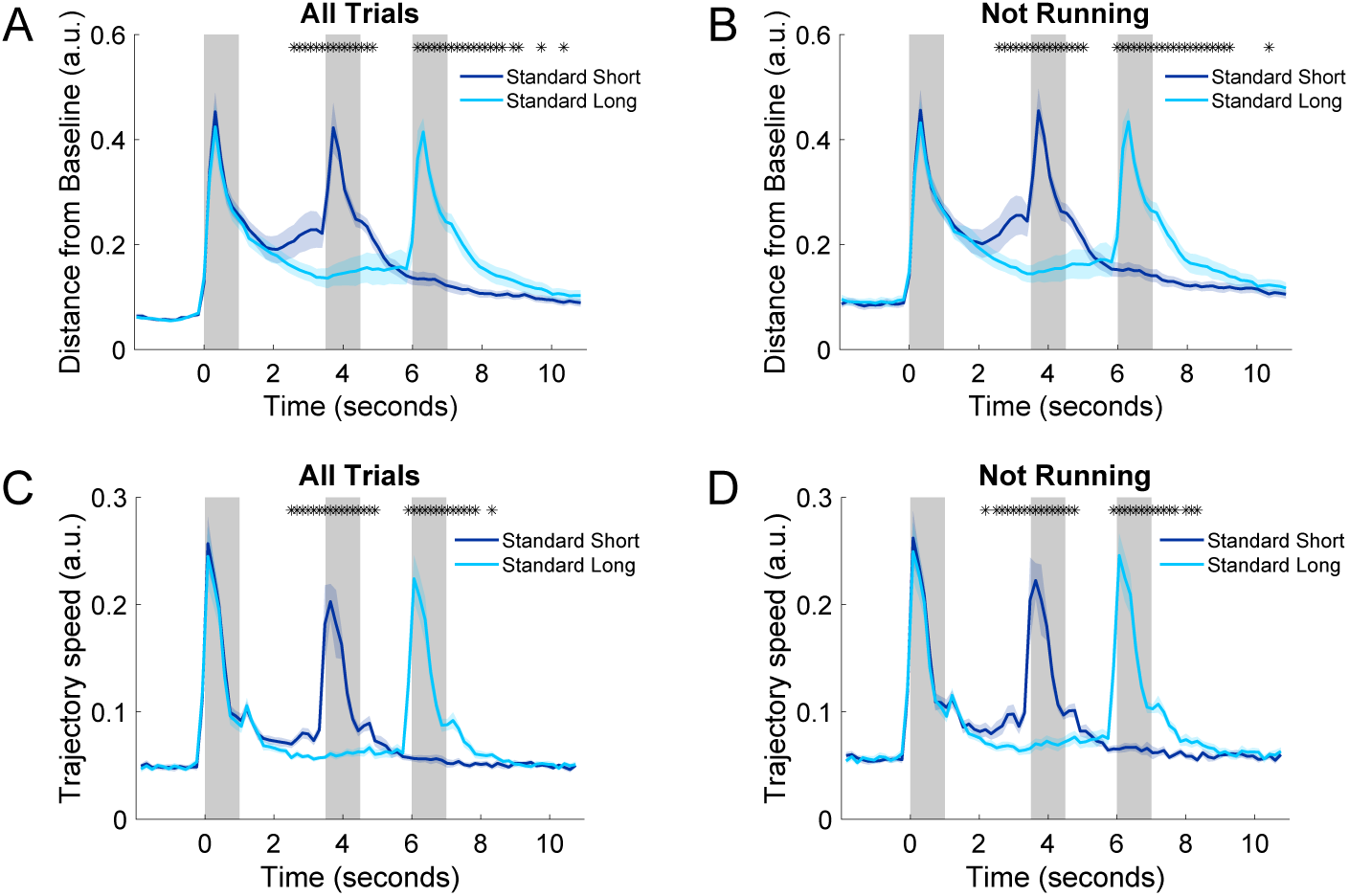
Locomotion running patterns did not explain effects of neural trajectories. **A)** Same as Figure 3 B-C but comparing standard short and standard long trials. **B)** Same as (A) but excluding trials in which a 1-second period of running occurred during the delay period. **C)** Same as Figure 3 D-E but comparing standard short and standard long trials. **D)** Same as (C) but excluding running trials.

